# RIP140 regulates HES1 oscillatory expression and mitogenic activity in colon cancer cells

**DOI:** 10.1101/2023.05.13.539637

**Authors:** Nour Sfeir, Marilyn Kajdan, Stéphan Jalaguier, Sandrine Bonnet, Catherine Teyssier, Samuel Pyrdziak, Rong Yuan, Emilie Bousquet, Antonio Maraver, Florence Bernex, Nelly Pirot, Florence Boissiere, Audrey Castet-Nicolas, Marion Lapierre, Vincent Cavaillès

## Abstract

**Background:** The transcription factor RIP140 (Receptor Interacting Protein of 140 kDa) regulates intestinal homeostasis and tumorigenesis through the Wnt signaling. In this study, we have investigated its effect on the Notch/HES1 signaling pathway.

**Methods:** The impact on HES1 expression and activity was evaluated in colorectal cancer (CRC) cell lines and in transgenic mice, invalidated or not for the *Rip140* gene in the intestinal epithelium. A tumor microarray and transcriptomic data sets were used to investigate RIP140 and HES1 expression in relation with patient survival. Statistical comparisons were performed with Mann-Whitney or Kruskal-Wallis or *Chi2* tests.

**Results:** In CRC cells, RIP140 positively regulated *HES1* gene expression at the transcriptional level via an RBPJ/NICD-mediated mechanism. In support of these *in vitro* data, RIP140 and HES1 expression significantly correlated in mouse intestine and in a cohort of CRC samples, analyzed by immunohistochemistry, thus supporting the positive regulation of *HES1* gene expression by RIP140.

Interestingly, when the Notch pathway is fully activated, RIP140 exerted a strong inhibition of *HES1* gene transcription controlled by the level of HES1 itself. Moreover, RIP140 directly interacts with HES1 and reversed its mitogenic activity in human CRC cells. In line with this observation, HES1 levels were associated with a better patient survival only when tumors expressed high levels of RIP140.

**Conclusions:** Our data identify RIP140 as a key regulator of the Notch/HES1 signaling pathway with a dual effect on *HES1* gene expression at the transcriptional level and a strong impact on colon cancer cell proliferation.

## BACKGROUND

Colorectal cancer (CRC) is the third leading cause of cancer deaths with a mortality rate of 33% in developed countries (1). Different genetic variations are involved in the development and progression of CRC (2). In addition, various signaling pathways which regulate intestinal development and differentiation play important roles in early stages of colorectal carcinogenesis (3),(4).

Increasing evidence has shown that the Notch signaling pathway controls many aspects of intestinal epithelium development and renewal (5),(6). The Notch pathway is an intercellular communication mechanism activated following the binding of a ligand of the jagged and delta family to a membrane receptor of the Notch family carried by the neighboring cell (7). The interaction between a Notch receptor and its ligand leads to a conformational change of the receptor at the origin of a double sequential cleavage. The intracellular domain of the Notch receptor (NICD for Notch intracellular domain) is translocated to the nucleus to induce the expression of target genes (such as HES 1-7 and HEY1-2) with the aid of the transcription factor RBPJ and coactivators such as MAML1 (Mastermind-like). In absence of Notch activation, RBPJ acts as a transcriptional repressor by recruiting a complex of co-repressors which comprises SHARP (SMRT and HDAC associated repressor protein), CtBP (C-terminal binding protein) and CtIP (CtBP interacting protein) (8).

The Notch signaling pathway plays an essential role in intestinal development and homeostasis by maintaining intestinal progenitors and stem cells in a state of proliferation while promoting their differentiation into absorptive cell lineage in the detriment of a secretory lineage differentiation (9). In addition, the Notch pathway has an oncogenic potential in colon cancer (10). One of its target genes, HES1, is the major effector in the intestine and plays an important role in CRC by regulating key parameters involved in tumorigenesis, including cell proliferation and differentiation (11),(12). Interestingly, in order to avoid an aberrant activity of the Notch pathway, HES1 exerts a negative feedback loop on its own promoter leading to an autonomous oscillatory expression (13). Finally, the promoter of the *HES1* gene is not only regulated by the Notch pathway but also by the Wnt pathway due to the presence of two TCF binding sites near the RBPJ site (14).

RIP140 (Receptor Interacting Protein, 140 kDa) also known as NRIP1 (Nuclear Receptor-Interacting Protein 1), was initially characterized as a transcriptional repressor of nuclear hormone receptors (15),(16). We and other then identified RIP140 as a coregulator of various transcription factors, including E2F1 (17), HIFs (18) and NFK-B (19). The repressive activity of RIP140 involves several inhibitory domains interacting with histone deacetylases and CtBPs (20) and is controlled by different post-translational modifications (21). Using a mouse model lacking the *Rip140* gene, a wide range of physiological processes were shown to be regulated by RIP140, including female fertility (22) and mammary gland morphogenesis (23), fat metabolism (24), proinflammatory cytokine response (25) and cognition (26).

In the intestinal epithelium, our laboratory demonstrated that RIP140 inhibited the Wnt/β-catenin signaling pathway and exerted an antiproliferative activity on CRC cells (27). Consequently, RIP140 expression decreased in CRC samples as compared to the adjacent healthy tissue. Interestingly, in sporadic CRC, RIP140 mRNA and protein levels significantly correlated with a better overall survival of patients and was identified as a good prognosis marker (27),(28). More recently, we demonstrated that RIP140 acts as a major regulator of SOX9 signaling with functional relevance in intestinal physiopathology, in relation with Paneth cell differentiation and colon cancer cell proliferation (29). In another recent study, we reported that RIP140 is involved in the regulation of microsatellite instability in CRC cells through the regulation of *MSH2* and *MSH6* gene expression (30). Interestingly, a frame shift mutation in the RIP140 coding sequence was identified in microsatellite instable CRCs with a familial history and this mutation appeared to be associated with intestinal tumorigenesis.

In the present work, we demonstrated that RIP140 strongly interferes with the Notch pathway by controlling *HES1* gene expression at the transcriptional level, participating in the HES1 regulatory negative feed-back loop, antagonizing HES1 mitogenic activity *in vitro* and impacting the prognostic value of HES1 in human CRC. Altogether, this work identifies RIP140 as a new regulator of the Notch/HES1 axis, which amplifies its role in the fine-tuning of intestinal tumorigenesis.

## MATERIALS AND METHODS

### Cell culture

The human colon adenocarcinoma cell lines SW620, LS174T, SW480, HT29 and HCT116 were cultured in RPMI, DMEM-F12 or McCoy’s medium supplemented with 10% FCS, 100U/ml penicillin, 100mg/ml streptomycin and 100mg/ml sodium pyruvate.

### Plasmids and treatments

pRL-TK-renilla and pGL3 plasmids were obtained from Promega (Charbonnieres, France). pEF-cmyc-RIP140 and pEGFP-RIP140 have been previously described (31). The reporter plasmids containing the firefly luciferase gene under the control of different fragments of the *RIP140* promoter were already described (32). The PCMV6-HES1 (33), PCMV5-NICD (34) and pRBPJ-Luc plasmids were kindly given by the corresponding laboratories. The lentiviral expression vectors EF.hHES1.Ubc.GFP (LV-HES1; #17624 (35)), EF.deltaBHES1.Ubc.GFP (DBD-HES1 DNA-binding domain mutant of HES1; #24982) were obtained from Addgene (Cambridge, USA). The reporter plasmids containing the firefly luciferase gene under the control of various fragments of the *Hes1* promoter (2.5 kb; #43806 (36)), (467bp; #41723 (37)) or 467bp lacking the RBPJ site; #43805) or the *Hes5* promoter (#41724 (37)) were obtained from Addgene (Cambridge, USA). Different fragments of the Hes1 promoter (WT or mutRBPJ) were PCR amplified and cloned into pGL3-basic plasmid previously digested with HindIII-XhoI or HindIII-KpnI to create I3, I1 and I1mutRBPJ reporter vectors which contained the -490/+46 or -125/+46 region of the RIP140 promoter with the wild-type sequence or with a mutation of the RBPJ site. All the engineered PCR constructs were sequenced. Sp1-Luc reporter and plasmids containing p21^WAF1/CIP1^ promoter (pWP101wild type) driving the luciferase reporter gene were provided by C. Seiser (Vienna BioCenter, Vienna, Austria) and Y. Sowa (Kyoto, Japan), respectively and as previously described (38). To inhibit the Notch pathway, cells were treated with 0.5 µM of an inhibitor of the gamma secretase, DBZ (Syncom, SIC 020042), for 24h.

### Luciferase and ChIP assays

Transient transfections with various promoter constructs were performed using Jet-PEI® (275ng of total DNA) according to the manufacturer instructions. SW620 and HT29 cells were seeded in 96-well plates (3×10^4^ and 1.5×10^4^ cells per well respectively) 24h prior to DNA transfection. Cells, in triplicates, were transfected with 25ng of firefly luciferase-based reporter constructs and 50ng renilla luciferase plasmid pRL. In cotransfection experiments, different amounts of expression plasmids were added. The pRL-TK-Renilla plasmid (Ozyme®) was used to normalize transfection efficiency. Firefly luciferase values were measured and normalized by the renilla luciferase activity. Values were expressed as relative luciferase activity (RLU) as mean ± S.D.

ChIP assays at the *HES1* promoter were performed in HT29 cells using the CHIP-IT kit (Active Motif, Carlsbad, USA). Sonicated chromatin was immunoprecipitated with antibodies against IgG (sc-3739, Santa Cruz Biotechnology, Inc, Heidelberg, Germany), H3pan (CC16310135, Diagenode, Liège, Belgium), and RIP140 (ab42126, Abcam, Paris, France). Immunoprecipitated DNA was amplified by qPCR using the primers listed in Supplementary Table S1.

### Small interfering RNA transfection

SW620 and HT29 cells were seeded in 6-well plates (8×10^5^ and 5×10^5^ cells per well respectively) 24h prior transfection. Small interfering RNA (siRNA) transfections targeting RIP140 (100pmol) or HES1 (200pmol) were performed using INTERFERin® transfection reagent. Cells were then incubated for 24h before RNA analysis by quantitative RT-qPCR.

### Real-time quantitative PCR (RT-qPCR)

Total RNA was extracted from cells using Quick-RNA kit (Zymo Research) according to the manufacturer’s instructions. RNA extraction from tissues or intestinal epithelial fraction was perform using EZNA® Total RNA kit (Omega Bio-tek) with bead column for more efficiency cell lyses. Total RNA (1µg) was subjected to reverse-transcription using qScript cDNA SuperMix (QuantaBio). RT-qPCR were performed with the Roche LightCycler® 480 instrument and the PerfeCTa SYBR Green FastMix (QuantaBio) and were carried out in a final volume of 10μl using 0.25µl of each primer (25μM), 5μl of the supplied enzyme mix, 2.5μl of H2O and 2μl of the template diluted at 1:6 (See Supplementary Table S1 for primer sequences). After pre-incubation at 95°C, runs corresponded to 35 cycles of 15s each at 95°C, 5s at 60°C and 15s at 72°C. Melting curves of the PCR products were analyzed using the LightCycler® software to exclude amplification of unspecific products. Results were normalized to RS9 or 28S housekeeping gene transcripts.

### Immunoblotting

RIPA solution was used to extract cell proteins. Cell extract were analyzed after migration of 50µg protein extract by Western blotting using a primary antibody against HES1 (#11988, Cell Signaling). Signals were revealed using a rabbit peroxidase-conjugated secondary antibody (1/5000, A6154 Sigma-Aldrich®) and a chemiluminescence substrate (ECL-RevelBlotPlus; GE Healthcare®) according to the manufacturer’s instructions. Protein quantifications were normalized with the β-actin signal (A3854; Sigma-Aldrich®).

### DuoLink proximity ligation assay

The proximity ligation assay was performed to visualize interactions using Duolink^®^ kit (Sigma-Aldrich®) according to the manufacturer instructions. SW620 or HT29 cells were plated on slides (5.10^4^ cells per well) 24h prior fixation with paraformaldehyde 3.7% and permeabilization with Triton X-100 1%. After blocking with BSA 1% for at least 3h, cells were incubated with two primary antibodies RIP140 (sc-9459, Santa-Cruz) and HES1 (#11988, Cell Signaling Technology) overnight at 4°C. A pair of oligonucleotide-labeled secondary rabbit and goat antibodies IgG (Duolink^®^ In Situ PLA^®^ Probes) was used according to the manufacturer’s instructions to bind to the primary antibodies. This pair of secondary antibodies generates a signal only when the two probes are in close proximity (40nm). The PLA signals were assigned using the Duolink^®^ In Situ Detection Reagents Orange (554nm laser line). Detection of the nucleus was done with the Hoechst (1/1000, Sigma-Aldrich®). Slides were counterstained with Hoechst (1/1000, Sigma Aldrich®) and mounted with Mowiol (Sigma-Aldrich®) for fluorescence microscopy. The images were obtained with ×40 magnification using an Axio Imager microscope (Zeiss).

### GST pull-down

Different fragments of RIP140 protein fused with GST: GST-RIP1(27-439), GST-RIP5(428-739), GST-RIP2(683-1158) and the GST(Control) were produced and purified as previously described (39). Unlabeled proteins were cell-free-synthesized using the TNT T7 Quick Reaction system according to the manufacturer instructions (Promega) and incubated with purified GST fusion proteins coupled with glutathione sepharose 4B fast flow beads (GE healthcare) in NETN buffer (20mM Tris PH8, 100mM NaCl, 1mM EDTA, 0,5 % Nonidet-P40, 1mM DTT, EDTA-Free protease inhibitors (Complete®) overnight at 4°C. After washing 4 times with NETN buffer, loading buffer 1X (50µl) was added to the beads and boiled for 5 min. Proteins were separated on a 10% SDS-PAGE.

### Histological and immunofluorescence analysis

Immunohistochemistry and immunofluorescence experiments were performed to detect RIP140 and HES1 protein expression. For Immunohistochemistry, mouse tissues were fixed with 3.7% paraformaldehyde, embedded in paraffin and sectioned (3μm). Following incubation in citrate buffer solution, immunohistochemistry analysis was performed using SignalStain® kit according to the manufacturer’s instructions. Paraffin-embedded tissue sections were first incubated in 3% hydrogen peroxide solution to block endogenous peroxidase activity. Each section was then incubated in blocking serum for 3h to reduce non-specific binding. Sections were then incubated with antibodies specific for RIP140 (Ab42126, Abcam) and HES1 (#11988, Cell Signaling) diluted in SignalStain® antibody diluent (1:100) overnight at 4°C. Incubation with labelled HRP anti-rabbit and visualization with diaminobenzidine as a substrate were performed. All slides were counterstained with haematoxylin and images were taken using NanoZoomer (Hamamatsu Photonics).

For immunofluorescence, cells were fixed with 3,7% paraformaldehyde and paraffin-embedded tissue sections were incubated in citrate buffer solution, then permeabilized with PBS-1% Triton for 30 minutes, blocked with PBS 1%BSA for at least 3 hours and incubated with the primary antibodies (RIP140, 1:100, ab42126; HES1, 1:100, #11988) overnight at 4°C, diluted in PBS 1%BSA. Revelation was performed using Alexa Fluor secondary rabbit antibodies IgG (AF488®, AF546®, 1/400, Invitrogen®). For co-localization experiment, we used the primary antibody (RIP140, 1:100, sc-9459) and the revelation was made using Alexa Fluor secondary goat antibody IgG (AF488®). Slides were counterstained with Hoechst (1/1000, Sigma Aldrich®) and mounted with Mowiol (Sigma-Aldrich®) for fluorescence microscopy. Staining quantification was performed at x40 magnification using the AxioVision software (Carl Zeiss®).

### TMA construction

TMA was constructed with FFPE tumor samples collected in the frame of the Clinical and Biological Database BCBCOLON (Institut du Cancer de Montpellier - Val d’Aurelle, France, Clinical trial Identifier #NCT03976960). Adenomas, primary adenocarcinomas, and metastatic lesions were sampled as two cores of 1 mm diameter. All samples were chemonaive. Tumor samples were collected following French laws under the supervision of an investigator and declared to the French Ministry of Higher Education and Research (declaration number DC-2008–695). Study protocol has been approved by the French Ethics Committee: CPP (Comité de Protection des Personnes) Sud Méditerrannée III (Ref#2014.02.04) and by the local translational research committee (ICM-CORT-2018-28). We have complied with all relevant ethical regulations for work with human participants, and informed written consent was obtained for all patients.

### IHC on human samples

The RIP140 detection has been described previously (30).HES1 immunostaining of human FFPE CRC samples was performed on a Dako Autostainer using the rabbit polyclonal antibody D6P2U (Cell Signalling Technology). The PT-Link® system (Dako) was used for simultaneous deparaffinization and antigen demasking for 15 minutes at 95°C. Endogenous peroxidases and non-specific signal (Flex Peroxidase Block and Protein Block, Dako) were blocked at room temperature. Sections were incubated for 30 minutes with the rabbit anti-human HES1 antibody diluted 1:100 in a low background diluent (Dako). A linker rabbit was used to amplify the signal. After two rinses, slides were incubated for 20 minutes at room temperature with an anti-rabbit antibody coupled with horseradish peroxidase (HRP) (Flex, Dako). Finally, 3,3’-Diaminobenzidine was used as a substrate for HRP and after rinsing, the slides were counterstained with hematoxylin, dehydrated and mounted with a coverslip.

Two readers performed independently the semi-quantitative analysis of immunohistochemical signals. The variations in HES1 or RIP140 expression observed between the different tumors allowed us to establish a score grid (no labeling: 0; weak labeling: 1; moderate labeling: 2; strong labeling: 3). For each sampled spot, the percentage of cells labeled with each intensity was reported. The overall expression was then calculated using the H-Score^2^ method (3 x % of cells with 3+ labeling intensity 2 x % of cells with 2+ intensity and 1 x % of cells with 1+ labeling intensity). The scores obtained thus varied from 0 to 300.

### Cell proliferation

Transient transfected cells were seeded in 96-well plates (6 replicates per condition), at a density of 5.10^3^ cells per well. After 4 days, 0.5mg/ml of 3-(4,5-dimethylthiazol-2-yl)-2,5-diphenyltetrazolium bromide (MTT) (Sigma-Aldrich®, St Louis, MO, USA) was added and incubated at 37°C for 3h. Formazan crystals were solubilized in DMSO and absorbance read at 560 nm on a spectrophotometer. Results were normalized to the mean optical density of the control.

Transiently transfected HT29 cells in the absence or presence of the siRNA targeting RIP140 were seeded at a density of 2500 cells/well into E-Plate 16 (ACEA Biosciences, Inc., San Diego, CA) containing 150μl per well of medium supplemented with 10% FCS. Dynamic monitoring of cell growth was determined every 12 hours during 6 days using the impedance-based xCELLigence system (Roche Applied Science, Germany). The cell index was derived from measured cell-electrode impedance that correlates with the number of viable cells.

### Animals

To generate the C57BL/6J mice line with conditional KO of the *Rip140* gene in the intestinal epithelium, RIPcKO^Int^ transgenic mice (obtained from the Yuan’s lab, Springfield, USA) were crossed with a mouse line expressing the CRE recombinase under the control of the villin promoter (40) and with the tumor-prone *Apc*^flox^ mouse strain (Jackson Laboratory). Animals were genotyped by PCR using ComR1 primers specific to the floxed region (see Supplementary Table S1 for primer sequences). All animals were maintained under standard conditions, on a 12:12-h light/dark schedule and fed chow diet *ad libitum*, according to European Union guidelines for use of laboratory animals. *In vivo* experiments were performed in compliance with the French guidelines for experimental animal studies (agreement D3417227).

### Statistical analysis

All experiments were conducted independently at least three times. Results were expressed as the mean ± standard deviation (S.D.). Statistical comparisons were performed with Mann-Whitney or Kruskal-Wallis tests. A probability level (*p* value) of 0.05 was chosen for statistical significance.

## RESULTS

### RIP140 increases *Hes1* gene expression *via* an RBPJ/NICD-mediated mechanism

To decipher the role of RIP140 on the Notch/HES1 pathway, we first measured its effect on *HES1* gene expression in different CRC cell lines in normal culture conditions. As shown in Figure 1A, ectopic expression of RIP140 in SW620 CRC cells significantly increased the level of *Hes1* mRNA, whereas the opposite effect was observed upon *Rip140* knock-down after transfection of a specific siRNA. The positive effect of RIP140 was also noticed on HES1 protein levels assessed by western-blot analysis (Figure 1B). Moreover, the same regulation of *Hes1* mRNA and protein levels was obtained in HT29 CRC cells (Supplementary Figure S1A and B).

**Figure 1:**
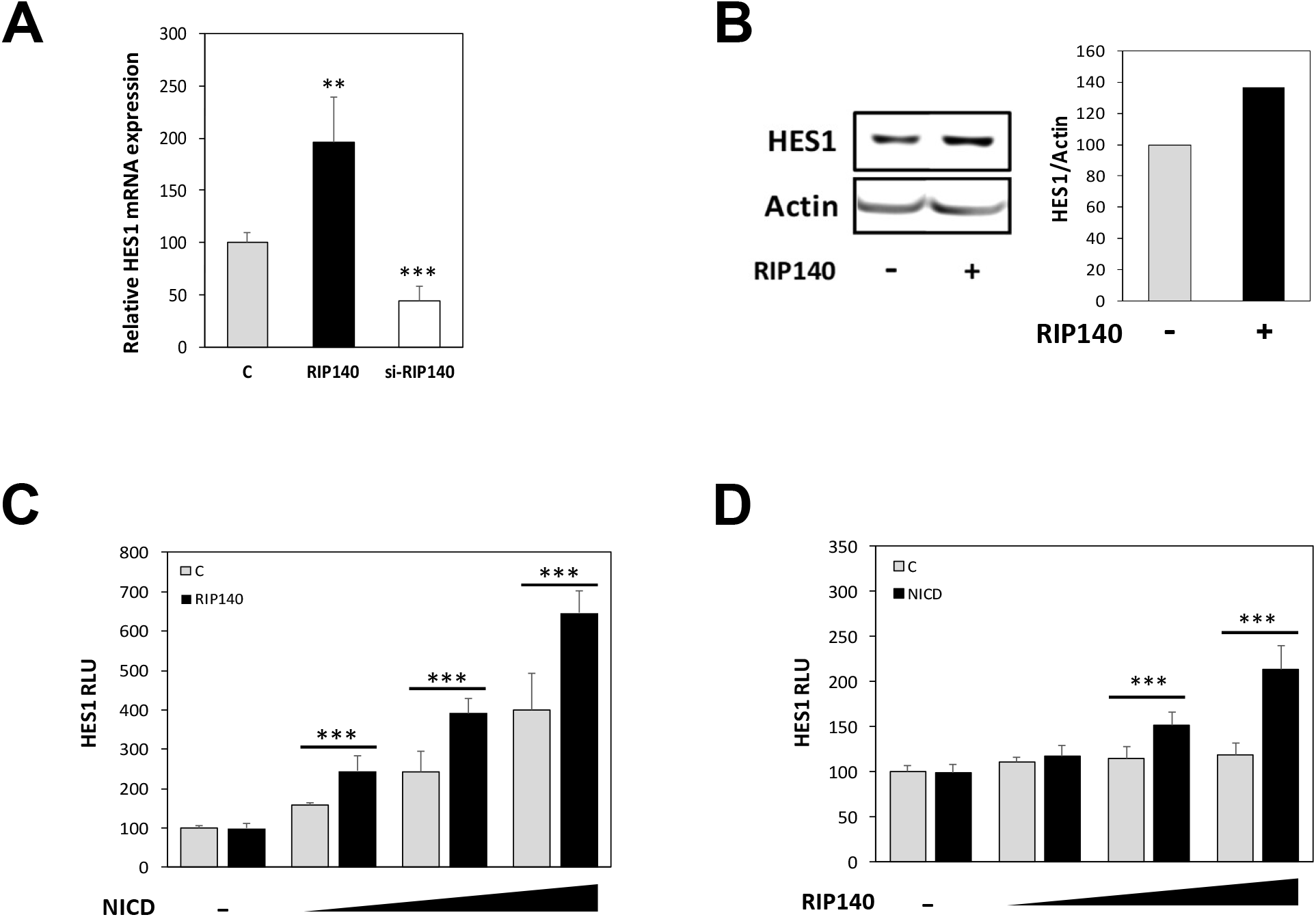
RIP140 increases HES1 expression at the transcriptional level. **(A)** *HES1* mRNA level in SW620 CRC cells transiently transfected with pEGFP or a control siRNA (C), pEGFP-RIP140 expression vectors (RIP140) or a RIP140 targeting siRNA (si-RIP140). Results are expressed as fold change ± S.D. relatively to controls; n = 3 to 5 independent experiments for each condition. **(B)** Western blot analysis of HES1 protein level in SW620 cells transiently transfected or not with a RIP140 expression vector. **(C)** Luciferase reporter assay performed with the reporter construct encompassing the 2.5 kb promoter region of *HES1* gene was transiently co-transfected into SW620 CRC cells with increasing doses of NICD expression vector in the presence (RIP140) or not (C) of the RIP140 expression vector. Relative luciferase activity was expressed as mean ± SD; n = 3 independent experiments. **(D)** A reporter construct encompassing the promoter region of the *HES1* gene (0.47kb) was transiently co-transfected into SW620 cells with increasing doses of RIP140 expression vector in the presence or not of NICD expression vector; n=3 independent experiments.

To further investigate the mechanisms involved in this regulation, we transiently transfected a luciferase reporter vector containing the promoter region of the *HES1* gene (encompassing the sequence from -2615 to + 46) (36) together with increasing doses of RIP140 expression vector. As shown in Figure 1C, we observed a dose-dependent transactivation by RIP140. In addition, we performed the same experiment in the presence or absence of a NICD expressing plasmid (Figure 1D) and found that the transactivation induced by the NICD-encoding plasmid was significantly increased by in the presence of the RIP140 expression vector. Altogether, these data clearly demonstrated that RIP140 exerted a NICD-dependent positive transcriptional regulation of *HES1* gene expression in human CRC cells.

To decipher further the underlying mechanisms, we generated several mutant versions of the *HES1* luciferase reporter construct. As shown in Figure 2A, the I1 plasmid which contains only the proximal promoter region of the *HES1* gene (- 125 to + 46) supported a positive NICD-mediated effect of RIP140. This NICD-dependent positive transcriptional regulation of the *HES1* gene by RIP140 was totally abolished upon mutagenesis of the RBPJ binding site, thus further demonstrating that the effect of RIP140 on *HES1* gene expression was RBPJ/NICD-mediated. This was confirmed by the use of an artificial reporter construct, which contains several RBPJ response elements upstream the minimal promoter driving the luciferase. Using this simplified RBPJ-Luciferase reporter plasmid, we obtained a significant stronger amplification of NICD-mediated transactivation by RIP140 both in SW620 (Figure 2B) and in HT29 CRC cells. Moreover, as above-mentioned for the *HES1* gene reporter (Figure 1D), the dose-dependent positive effect of RIP140 on the RBPJ-Luciferase reporter plasmid was observed only in the presence of NICD (Figure 2C). Finally, in accordance with these observations, we validate the NICD-mediated positive effect of RIP140 on *HES4* mRNA expression (Figure 2D) and on *HES5* gene transcription (Supplementary Figure S1C), both genes being also Notch target genes (41),(42). Altogether, these data demonstrated that, in human CRC cells, RIP140 increases the Notch pathway as demonstrated by the induction of RBPJ/NICD-mediated transcription of several targets of the Notch signaling, including the *HES1* gene.

**Figure 2:**
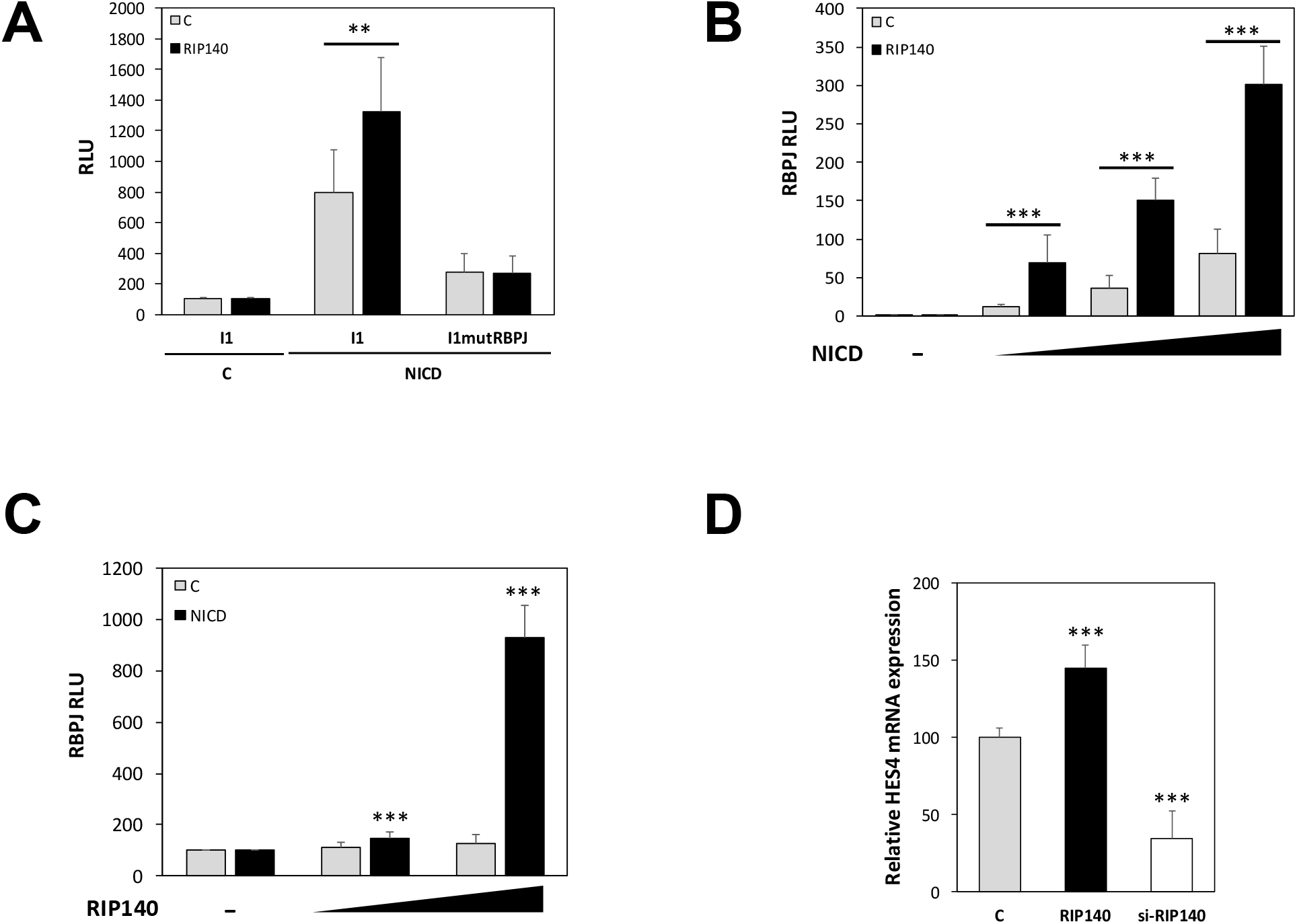
RIP140 regulates HES1 *via* an RBPJ/NICD mechanism. **(A)** The very proximal promoter region of HES1 with the RBPJ binding site and a remaining HES1 binding site (I1 construct) and I1 lacking the RBPJ site (I1mutRBPJ construct) were transiently co-transfected into SW620 cells with doses of NICD and RIP140 expression vectors. Results are expressed as fold change ± S.D. relatively to controls; n = 3 independent experiments. **(B)** Luciferase reporter assay performed on an artificial reporter construct containing 6 RBPJ-binding sites in the same conditions as in panel C of Figure 1. Relative luciferase activity was expressed as mean ± SD; n = 3 independent experiments. **(C)** Luciferase reporter assay performed on an artificial reporter construct containing 6 RBPJ-binding sites in the same conditions as in panel F; n=3 independent experiments. **(D)** *HES4* mRNA level in SW620 cells measured under the same condition as in panel A. Results are expressed as fold change ± S.D. relatively to controls; n=3 independent experiments for each condition.

### RIP140 is a HES1 target gene

As observed for other transcription factors engaged in negative feed-back loop with RIP140 including for instance the estrogen receptor (43) or the E2F1 transcription factor (44),(45), we expected a possible induction of the *RIP140* gene transcription by HES1. As shown in Figure 3A, HES1 ectopic expression indeed significantly increased the accumulation of the RIP140 protein detected by immunofluorescence. The same regulation of *RIP140* gene expression was observed at the mRNA level upon HES1 ectopic expression and the opposite effect was obtained upon HES1 knock-down using a specific siRNA in SW620 cells (Figure 3B) and similar results were obtained in HT29 cells (Supplementary Figure S2A).

**Figure 3:**
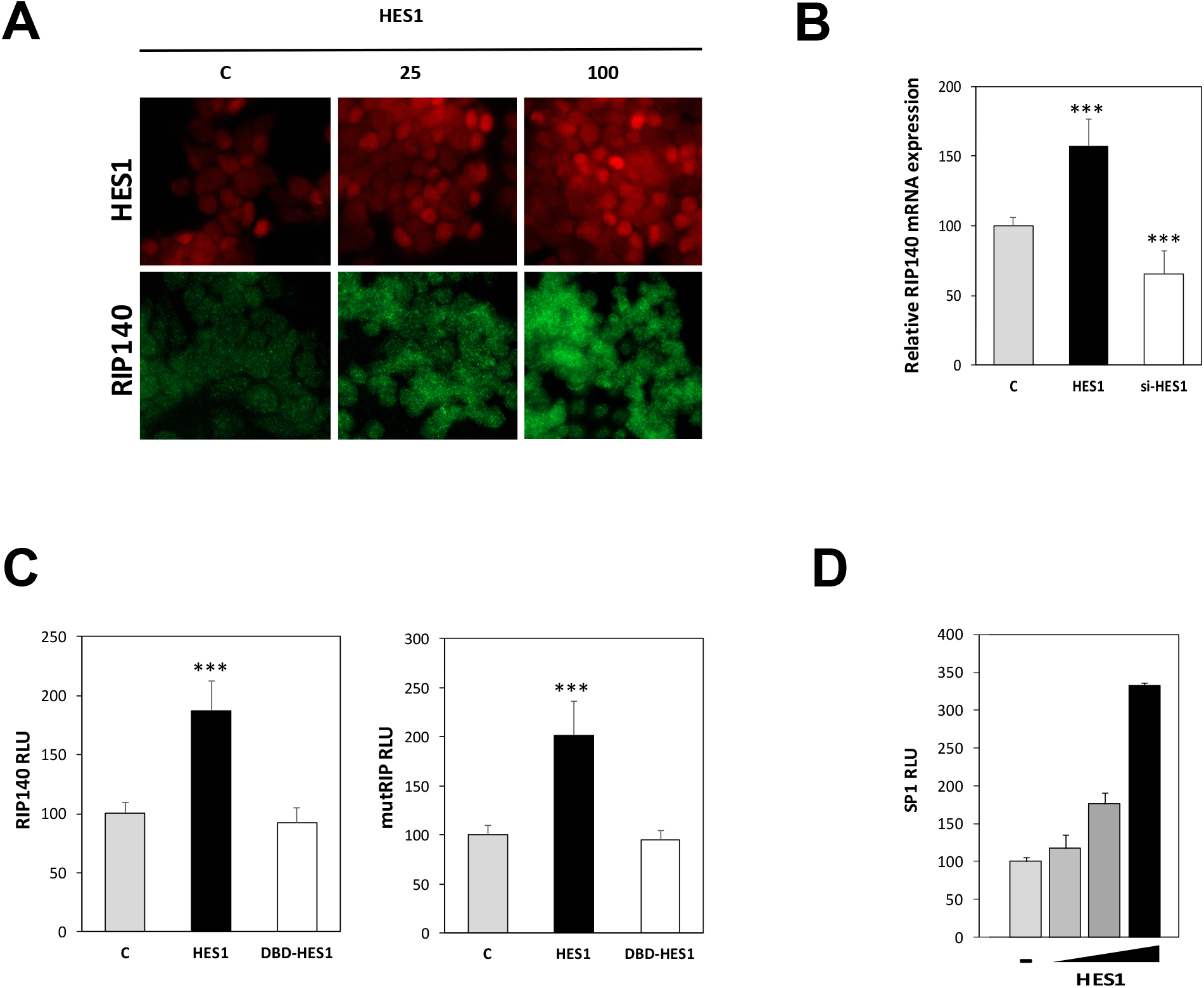
RIP140 gene expression is increased by HES1. **(A)** Immunofluorescence analysis (40x) of HES1 and RIP140 protein levels in SW620 cells transiently transfected with increasing doses of HES1 expression vector (25 and 100 ng). **(B)** *RIP140* mRNA level in SW620 cells transiently transfected or not with HES1 or with a siRNA targeting the *HES1* mRNA (si-HES1). Results are expressed as fold change ± S.D. relatively to the control; n = 4 to 6 independent experiments. **(C)** The same reporter constructs of panel D were transiently co-transfected into SW620 cells with HES1 or DBD-HES1 expression vectors. Relative luciferase activity was expressed as mean ± SD; n = 3 independent experiments**. (D)** An artificial reporter construct encompassing three SP1 binding sites was transiently co-transfected into SW620 cells with increases doses of HES1 expression vectors.

Interestingly, this regulation of RIP140 expression by the HES1 pathway was noticed at the transcriptional level upon HES1 ectopic expression (Figure 3C). These data were obtained on a luciferase reporter construct containing either the promoter region of the human *RIP140* gene encompassing the sequence from -814 to +106 (left panels) or a more proximal sequence from -158 to + 106 (right panels) (32). As expected, the DBD mutated version of HES1 was unable to increase *RIP140* gene transcription. It should be noted that the same opposite effects were measured in HT29 CRC cells (Supplementary Figure S2C and D) and observed, at the transcriptional level, on the murine *Rip140* promoter, in both SW620 and HT29 CRC cells (Supplementary Figure S2E). Moreover, a similar positive effect upon HES1 ectopic expression was observed on a reporter construct encompassing the luciferase gene controlled by three Sp1 binding sites(38) [(Figure 3D). This suggests that the regulation of *RIP140* gene expression by HES1 is possibly Sp1-mediated.

### Correlation between RIP140 and HES1 expression in mouse and human tissues

The above-mentioned molecular analyses performed in human cancer cell lines unraveled positive transcriptional regulatory links between RIP140 and HES1. In order to assess the relative expression of the two transcription factors in more physiological conditions, we analyzed their expression at the mRNA and protein levels in mouse tissue samples.

We used transgenic mice exhibiting a specific invalidation of the *Rip140* gene in the intestinal epithelium (RIPcKO^int^). These animals were obtained by crossing mice bearing the floxed *Rip140* gene (gift from R Yuan, Southern Illinois University, Springfield, USA) with a strain expressing the Cre recombinase under the control of the *Villin* gene promoter (40). As shown in Figure 4A and B, we observed a strong correlation between *Rip140* and *Hes1* mRNA levels (r=0.506; p<0.0005). The lowest levels of *Hes1* mRNA were measured in the intestinal epithelium of mice with the homozygous deletion of the *Rip140* gene as compared to wild-type mice (*Hes1* mRNA levels being intermediate in mice with heterozygous deletion of the *Rip140* gene). These results were validated at the protein level by immunohistochemistry in tissue sections of intestine from the RIPcKO^int^ compared to wild-type animals. The nuclear HES1 staining observed in the intestinal crypts was significantly decreased in the epithelium of RIPKO mice (Figure 4C). In line with this observation and our previous result, we also found a decrease of HES1 staining in the intestinal tumors developed by the RIPcKO^int^ mice crossed with the tumor-prone *Apc*^flox^ mouse strain, as compared to the control animals (Figure 4D).

**Figure 4:**
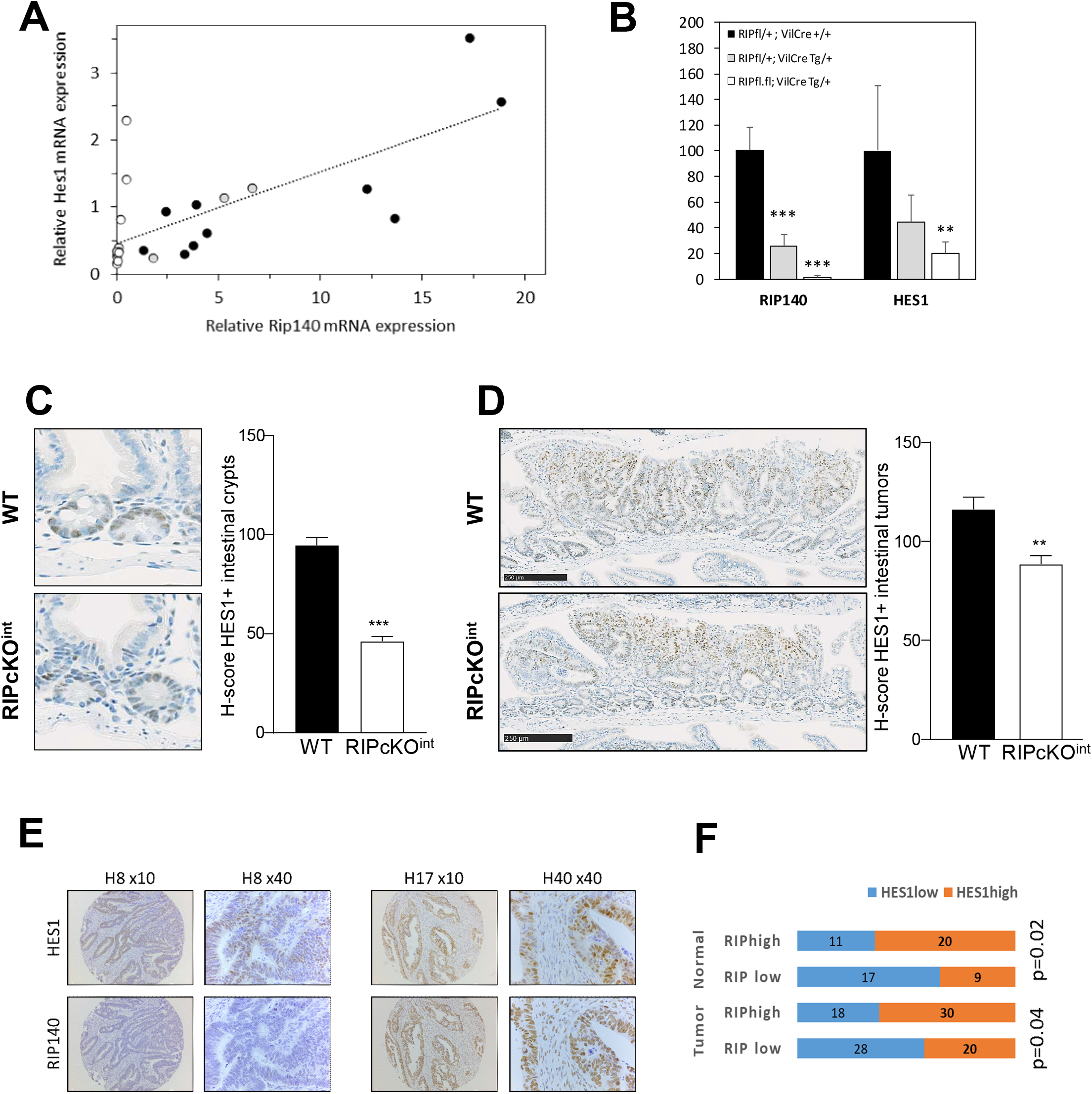
Positive correlation between RIP140 and HES1 expression in mouse intestine tissues and in human CRCs. **(A)** Correlation between *Rip140* and *Hes1* gene expression in intestinal epithelium cell fraction of conditional RIP140 KO mice (RIP^fl/+^; VilCre^Tg/+^ in grey and RIP^fl/fl^; VilCre^Tg/+^in white) compared to their wild-type littermates (RIPWT; in black). **(B)** *Rip140* and *Hes1* gene expression in intestinal epithelium cells fraction of *Rip140* conditional KO mice (RIP^fl/+^; VilCre^Tg/+^ and RIP^fl/fl^; VilCre^Tg/+^; RIPcKO^Int^) compared to their wild-type littermates (RIP^fl/+^; VilCre^+/+^; WT). **(C)** Immunohistochemistry and quantification of HES1 staining (H-Score values) in normal intestinal crypts of RIPcKO^Int^ mice compared to their wild-type littermates (WT) (40x). **(D)** Same as in **(C)** in intestinal tumors from RIPcKO^Int^/APC^fl/+^ mice compared to their littermates (WT). **(E)** Example of HES1 and RIP140 IHC staining performed as described in the Material and Method section and illustrating low (left panels) or high (right panels) intensity staining. The issue sample name and the tumor magnification are indicated above the photographs. **(F)** Schematic representation of the distribution of tumors with low or high HES1 expression (median cut-off) in the two subgroups of CRC with low (RIPlow) or high (RIPhigh) RIP140 expression (median cut-off). Statistical analysis was performed using the *Chi-2* test and p-values are indicated.

To validate these data in human tissues, we monitored the expression of RIP140 and HES1 proteins by immunohistochemistry in a cohort of human CRC biopsies. This cohort comprised 45 primary tumors of different stages (7 stage 0, 9 stage I, 10 stage II, 12 stage III, 7 stage IV) and 9 metastases (see Table 1 for patient characteristics). The expression of RIP140 and HES1 were quantified both in the non-tumoral mucosa and in the adenocarcinoma, as described in the Material and Methods section. As shown in Figure 5E and F, the nuclear staining of the two markers was significantly correlated in both normal and tumoral intestinal tissues.

**Table 1:**
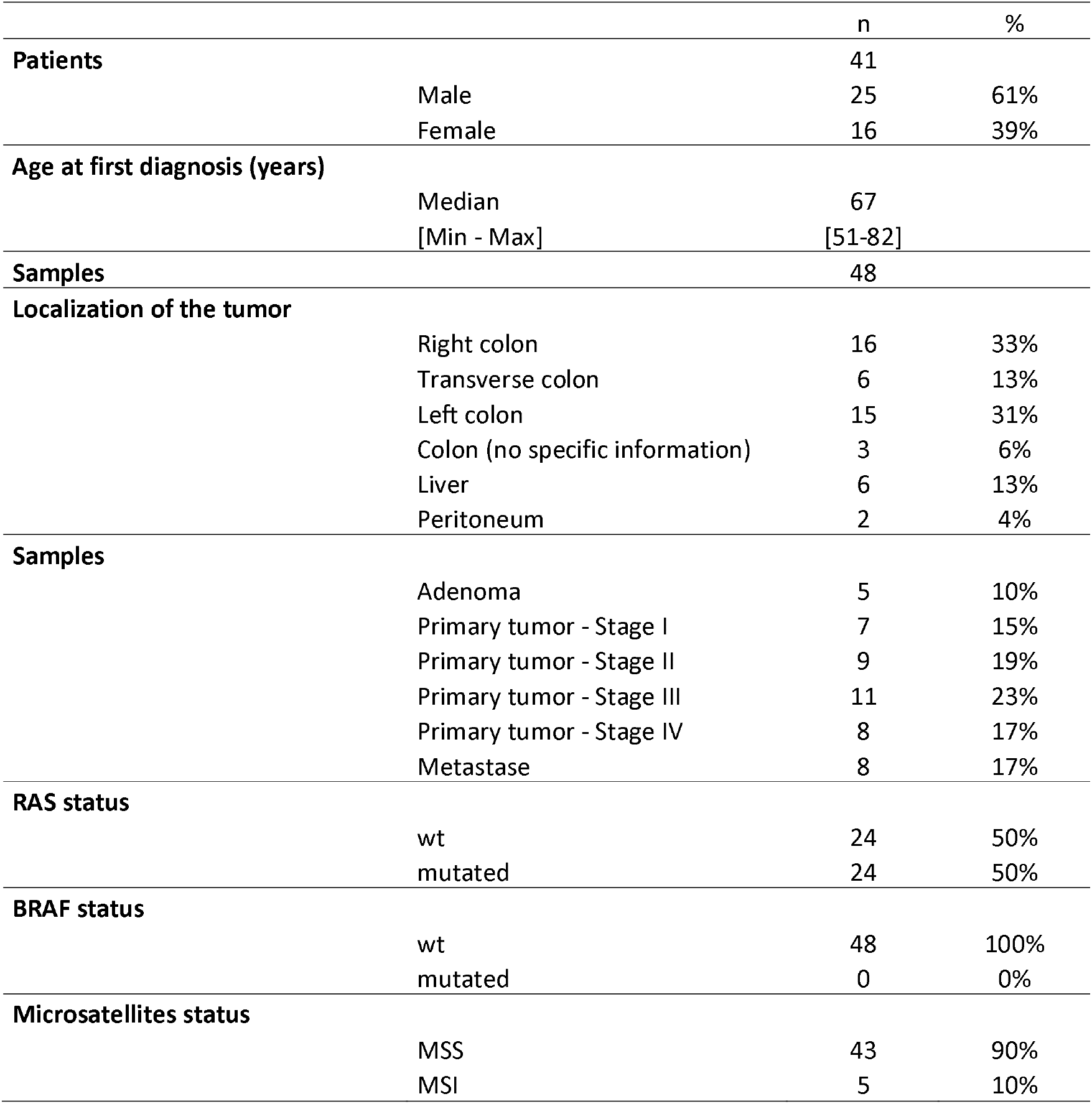
Characteristics of CRC patient cohort (n=45).

**Figure 5:**
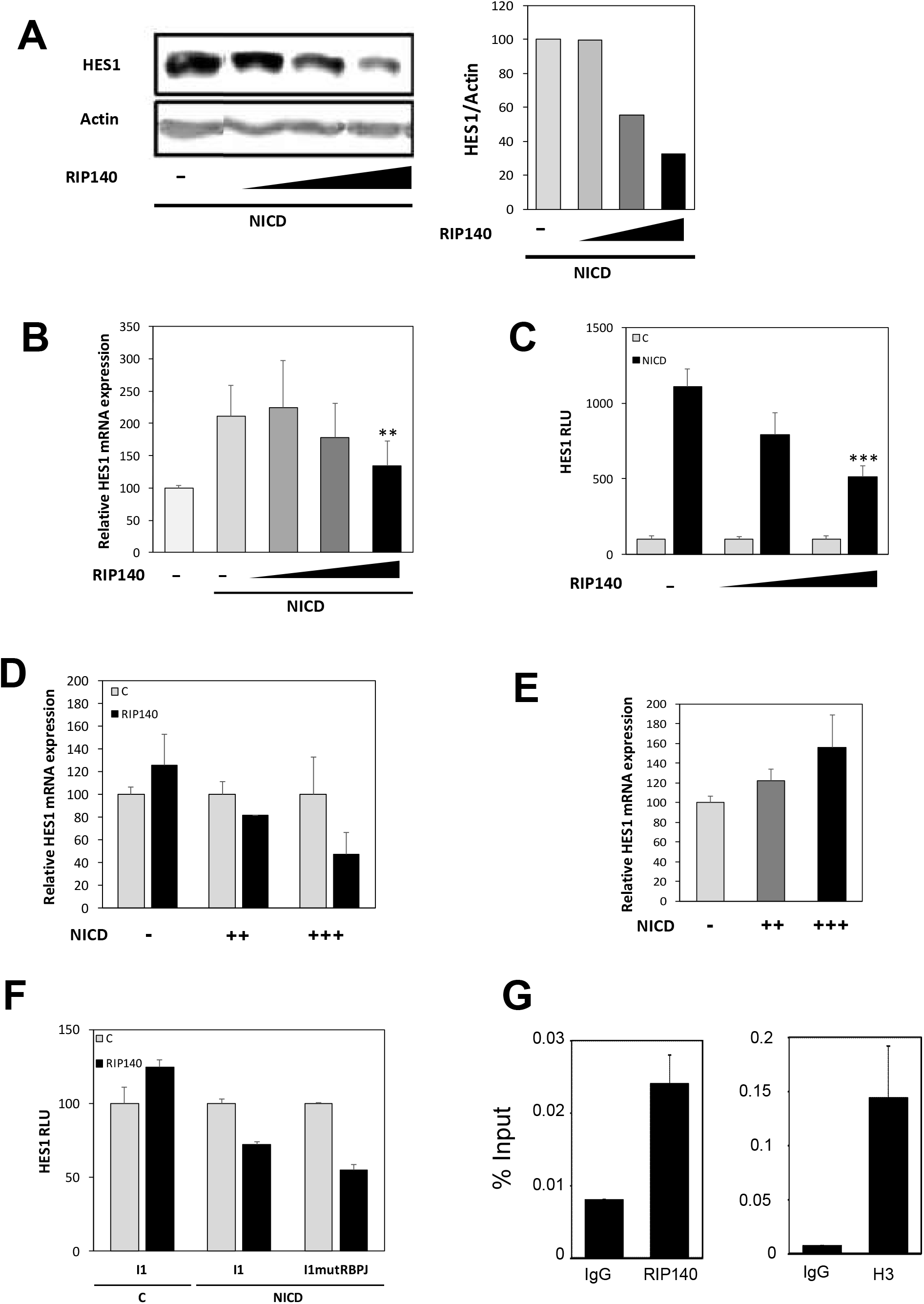
RIP140 inhibits HES1 gene expression when it is highly expressed. (**A)** Western blot analysis of HES1 protein level in SW620 cells transiently transfected with increasing doses of RIP140 expression vector (0,1, 0,5 and 1 µg) and a high dose of NICD expression vector (1 µg). (**B)** *HES1* mRNA level in SW620 cells measured under the same conditions as in panel A. Results are expressed as fold change ± S.D. relatively to the control n = 3 independent experiments. **(C)** Luciferase reporter assay performed on the *HES1* gene promoter (0.47kb) construct in SW620 cells transiently co-transfected with increasing doses of RIP140 expression vector in the presence or not of NICD expression vector. Relative luciferase activity was expressed as mean ± SD; n = 3 independent experiments **(D)** RT-qPCR analysis of *HES1* mRNA levels (top panel) in SW620 cells transiently transfected with increasing doses of NICD expression vector in the presence (RIP140) or not (C) of RIP140 expression vector. **(E)** Quantification of HES1 mRNA levels in the different NICD conditions shown in panel (D) without ectopic expression of RIP140. **(F)** I1 and I1mutRBPJ reporter constructs were transiently co-transfected into SW620 cells in the presence or not of NICD and/or RIP140 expression vectors. **(G)** ChIP assay using HT29 cells and anti-IgG, anti-RIP140 or anti-H3pan antibodies. Purified DNA was amplified by qPCR using HES1 promoter primer pairs (see upper panel in E). For all panels: * p < 0.05, ** p < 0.01 and *** p < 0.001 (Mann–Whitney test).

### Repressive effect of RIP140 when HES1 is expressed at high levels

As described above, our data evidenced a clear positive transcriptional regulation of the *HES1* gene by RIP140. However, the role of RIP140 in the control of *HES1* gene expression appeared more complex since we noticed in a reproducible way that, in conditions where HES1 expression was strongly induced by NICD, RIP140 exerted an inhibition of *HES1* gene transcription. Indeed, in such conditions, RIP140 behaved clearly as an inhibitor of *HES1* gene expression, as shown at the protein (Figure 5A), mRNA (Figure 5B) and transcriptional level (Figure 5C). Again, similar results were obtained in HT29 CRC cells (Supplementary Figure S3A and B) and on the *HES4* mRNA expression (Supplementary Figure S3C) thus demonstrating that the negative regulation exerted by RIP140, when the Notch pathway was fully activated, was a common effect.

As shown in Figure 5D, the switch in the effect of RIP140 from a positive to a negative regulation was clearly associated with the NICD induction of *HES1* gene expression (Figure 5E). The same switch from an activator to a repressor, according to NICD ectopic expression, was observed in HT29 cells at the transcriptional level on the *HES1* reporter construct (Supplementary Figure S3D). The inhibitory effect of RIP140 was also detected on the I1 construct encompassing the *HES1* proximal promoter (Figure 5F). More importantly, the negative effect of RIP140, which occurred at high NICD levels, was not abolished by the mutation of the RBPJ binding site (Figure 5F).

Finally, ChIP experiments (Figure 5G) demonstrated the specific recruitment of NRIP1 to the *HES1* promoter sequence that encompassed the RBPJ and HES1 binding sites that conferred regulation by NRIP1 in the above-mentioned luciferase assays. The amplification signal was lower than that observed after anti-histone H3 ChIP (H3) but higher than with control IgG. Moreover, we did not observe any recruitment to an irrelevant region.

Collectively, these data strongly suggested that, depending on HES1 protein level, RIP140 behaved as a direct activator or a repressor of *HES1* gene transcription.

### RIP140 is involved in the HES1 negative transcription feed-back loop

The oscillatory expression of the *HES1* gene and its regulation by a negative feedback loop have been reported more than fifteen years ago (46). One hypothesis that might recapitulate the effects of RIP140 on *HES1* gene expression relies on the participation of RIP140 in this HES1 negative feed-back loop.

To validate this hypothesis, we set up an assay allowing the quantification of the endogenous *HES1* gene expression in the presence or not of ectopic expression of the HES1 and/or RIP140 proteins. As shown in Supplementary Figure S4A (top scheme), we designed a specific RT-qPCR based quantification of the endogenous *HES1* gene expression by using specific primers in the 3’UTR region of the HES1 mRNA. Upon HES1 ectopic expression monitored using primers located in the coding sequence of the gene and allowing quantification of both ectopic and endogenous HES1 mRNAs (Supplementary Figure S4A - left panel), we observed the previously described negative feed-back loop (46) leading to a decreased endogenous HES1 RNA level (right panel).

Using this assay, we confirmed that the negative effect of RIP140 on endogenous *HES1* mRNA levels was indeed dependent on HES1 ectopic expression (Figure 6A). More importantly, we demonstrated that RIP140 was required for the HES1 negative feed-back-loop since the negative effect of HES1 on its own expression was abolished (and even reversed) upon siRNA silencing of *RIP140* gene expression (Figure 6B). The reinforcement by RIP140 of the inhibitory effect of HES1 on its own expression was also observed at the transcriptional level using the I3mutRBPJ reporter construct containing the HES1 promoter region mutated on the RBPJ binding sites and thus bearing only the HES1 response elements (Figure 6C). The importance of RIP140 in the inhibitory effect of HES1 was confirmed at the transcriptional level upon siRNA silencing of *RIP140* gene expression (Figure 6C). Finally, these results were strengthened by data showing that ectopic expression of RIP140 in SW620 cells increased the oscillatory expression of the HES1 mRNA as measured by immunofluorescence (Figure 6D).

**Figure 6:**
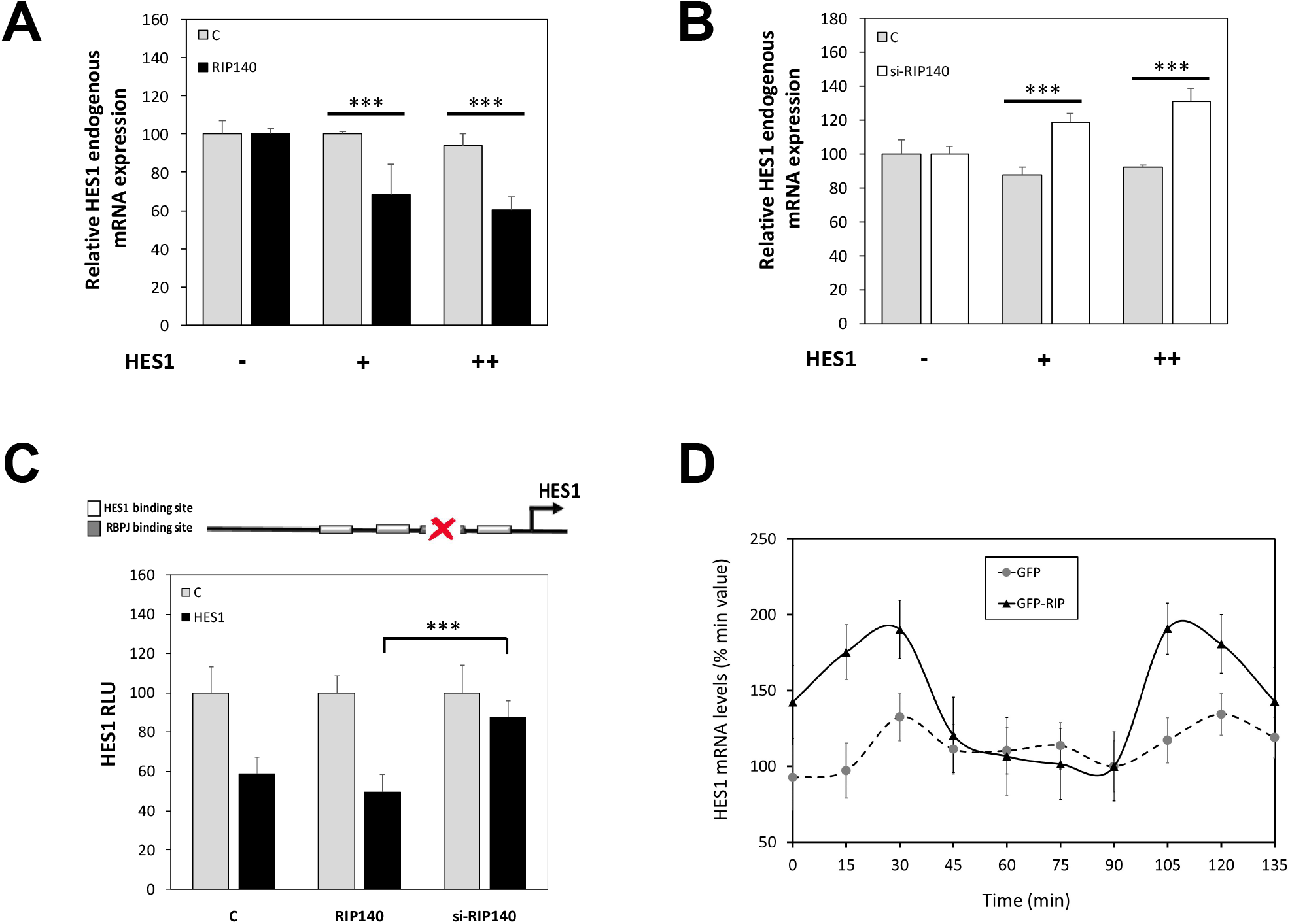
RIP140 is involved in the HES1 feed-back loop. **(A)** *HES1* endogenous gene expression in SW620 cells transiently transfected with increasing doses of HES1 expression vector in the presence or not of RIP140 expression vector. **(B)** *HES1* endogenous gene expression in SW620 cells transiently co-transfected with increased doses of HES1 expression vector together with a control siRNA (C) or a siRNA targeting RIP140 (si-RIP140). Results of panel B are expressed as fold change ± S.D. relatively to the control; n = 3 independent experiments. **(C)** The *HES1* gene promoter construct (I3mutRBPJ construct) containing the three HES1 binding sites and the mutated RBPJ binding site was transiently co-transfected in SW620 cells. The RIP140 expression vector or a siRNA targeting RIP140 (si-RIP140) together with the HES1 expression vector was co-transfected or not; n = 3 independent experiments. (**D)** Oscillatory expression of HES1 mRNA in SW620 cells transiently transfected with pEGFP (GFP) or pEGFP-RIP140 expression vectors (GFP-RIP140). Results are expressed as fold change ± S.D. relatively to minimum value; n = 3 independent experiments for each condition.

To characterize further the effect of RIP140 on HES1 repressive activity, we investigated whether the two proteins were able to interact. As shown in Figure 7A, GST pull-down experiments clearly demonstrated that the N-terminal region (fragment RIP1 that encompasses the RIP140 sequence from amino acid 27 to 439) was able to bind to the *in vitro* translated HES1 protein. The interaction between the two endogenous proteins was confirmed by proximity ligation assay (Figure 7B) and co-localization experiment in intact SW620 CRC cells (Figure 7C). Similar results were obtained in HT29 CRC cells (Supplementary Figure S4C-D). Altogether, these data supported the conclusion that RIP140 interacts with HES1 and is required for its full repressive activity, in particular on its own promoter.

**Figure 7:**
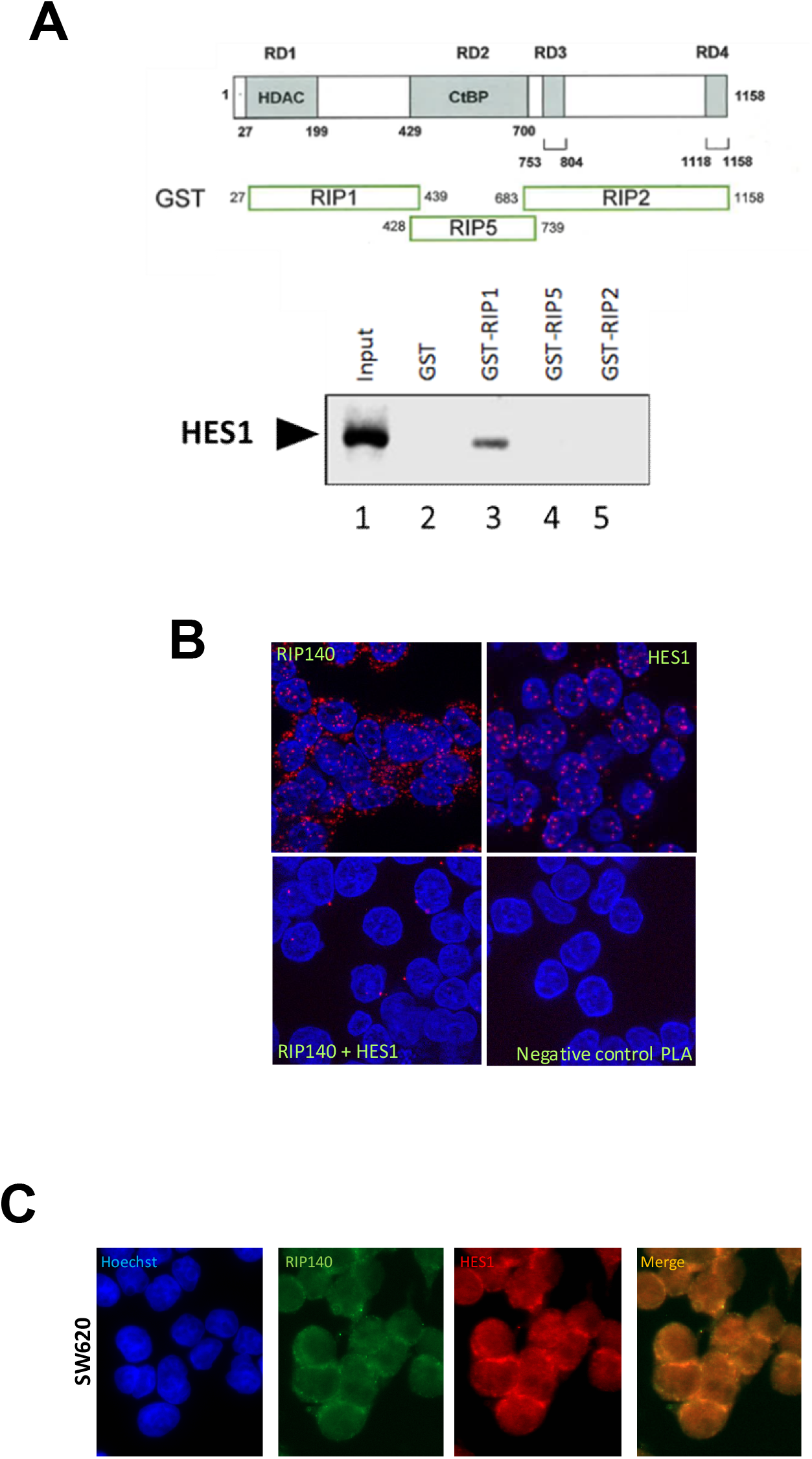
RIP140 interacts with HES1. **(A)** Analysis of RIP140 *in vitro* interaction with HES1 protein by GST pull down. The HES1 protein input is shown in lane1. The figure is representative of two independent experiments. **(B)** DuoLink proximity ligation assay performed to visualized endogenous HES1 and RIP140 interaction in SW620 cells. **(C)** Double immunofluorescence analysis (40x) of HES1 and RIP140 protein levels in SW620 cells.

### RIP140 switches the mitogenic effect of HES1 on CRC cell proliferation

We then investigated whether this intimate cross-talk between the two transcription factors might be relevant in CRC pathogenesis. We first performed Kaplan-Meier analyses of CRC patient survival based on HES1 expression. In the whole cohort, HES1 levels were not significantly associated with patient survival (Supplementary Figure S5A). Same results were obtained when we analyzed the tumors with the lowest level of RIP140 based on median cut-off (Figure 8A). By contrast, as shown in Figure 8B, high level of HES1 and RIP140 was associated with an increased patient survival.

**Figure 8:**
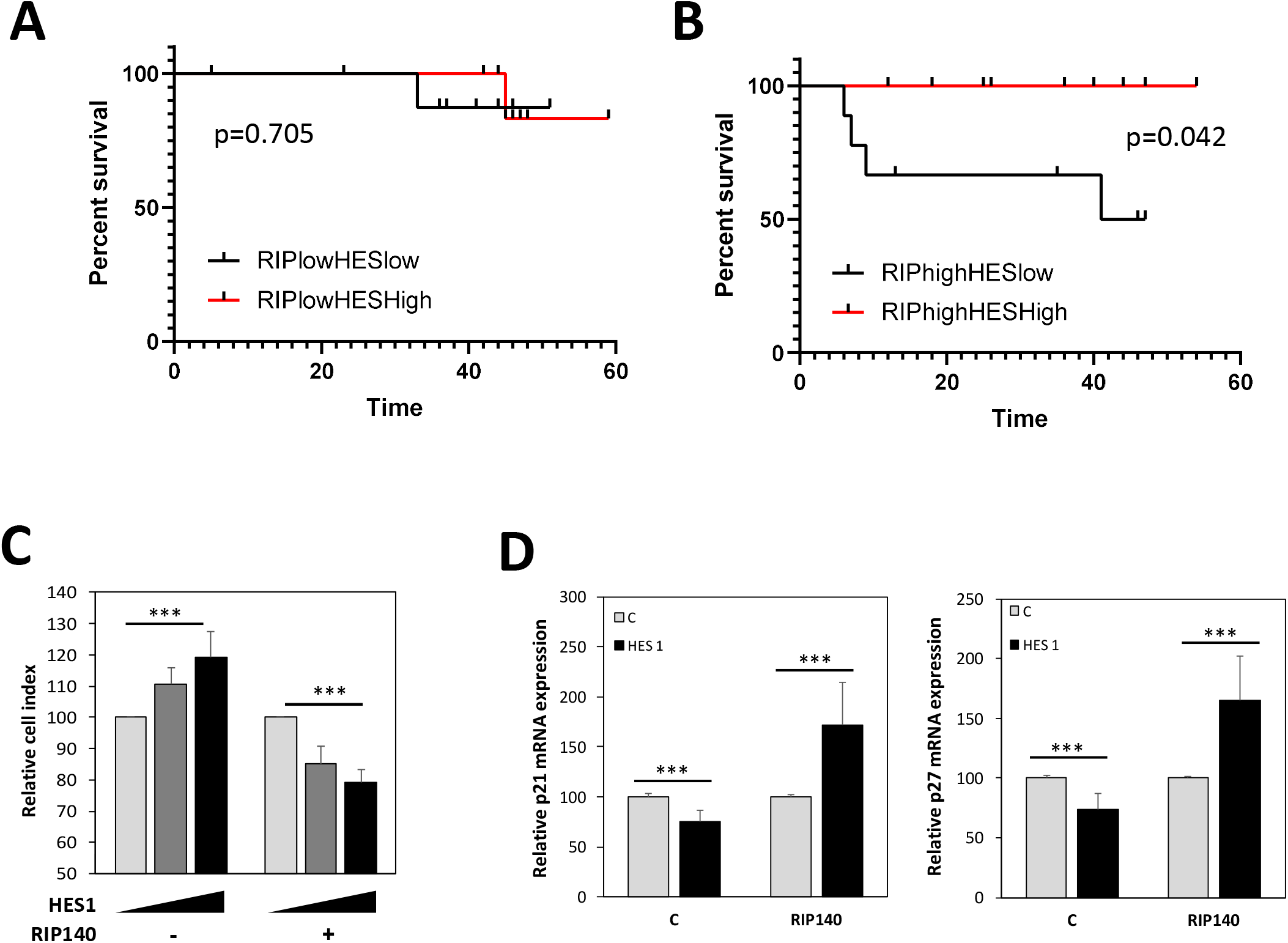
Crosstalk between RIP140 and HES1 in human CRC. **(A)** and **(B)** Kaplan-Meier analysis of the cumulative OS of patients with low or high HES1 staining IRS in their tumors was performed on the groups exhibiting low (panel A) or high (panel B) RIP140 staining IRS (best cut-off threshold). A log-rank test was used for statistical analysis**. (C)** The proliferation of SW620 cells transiently transfected with increasing doses of HES1 expression vector in the presence or not of RIP140 expression vector was quantified at day 4 using an MTT assay. Results represent fold change ± S.D. *vs* levels in control cells; n=2 independent experiments. **(D)** *p21* (left panel) and *p27* (right panel) mRNA levels in HT29 cells transiently transfected with RIP140 and HES1 expression vectors. Results are expressed as fold change ± S.D. relatively to control; n = 3 independent experiments.

These IHC data were confirmed at the level of mRNA expression by reanalyzing transcriptomic data from the TCGA (http://tcga-data.nci.nih.gov/docs/publications/coadread_2012/) (47). Using the Kaplan-Meier Plotter database, we reanalyzed RNAseq data obtained on colon adenocarcinoma from 452 patients. This cohort was separated into two groups of 226 patients with low and high RIP140 expression in the corresponding tumors using the median as a cutoff value (Supplementary Figure S5B and C, respectively). As observed by IHC, we confirmed a statistically significant association of high expression of HES1 with a decreased risk of death in colon cancer patients, only when their tumor express high RIP140 expression (Supplementary Figure S5C, p=0.048). Altogether, the data strongly suggested that RIP140 influences the biological activity of the HES1 protein, in accordance with our data demonstrating a strong effect of RIP140 on HES1 expression and transcriptional activity.

We then attempted to demonstrate experimentally this impact of RIP140 on HES1 oncogenic activity. As shown in Supplementary Figure S5D, we first validated that RIP140 exerted a clear antiproliferative activity in CRC cells since its knockdown in human CRC cells produced a significant mitogenic effect. These data confirmed previous results obtained in the HCT116 cell line (27) and were validated in other CRC cell lines. Very interestingly, when we tested the simultaneous ectopic expression of HES1 and RIP140, we clearly observed that overexpression of RIP140 switched HES1 effect from a stimulation to an inhibition of CRC cell proliferation (Figure 8C. Similar results were obtained on the expression of the *p21* and *p27* genes (Figure 8D). These two cyclin-dependent kinase inhibitors are key regulators of cell proliferation and survival which induce cell cycle arrest by inhibiting the activity of several cyclin-dependent kinases (48). These data strengthened the major switch exerted by RIP140 on the regulation of CRC cell proliferation by HES1. Altogether, they also explain why HES1 has a good prognosis value in CRC with high levels of RIP140.

## DISCUSSION

Colorectal cancer (CRC) is a frequent neoplasm that implicates the deregulation of multiple signaling pathways involved in the control of intestinal epithelial cell differentiation and proliferation. We previously described that the transcription factor RIP140 was an important player in the regulation of intestinal homeostasis and tumorigenesis through the control of *APC* gene expression and activation of the Wnt pathway (27). In the present study, we demonstrated that RIP140 also finely regulates the Notch signaling pathway and strongly cross-talks with the HES1 transcription factor with an impact on colon cancer cell proliferation and CRC prognosis.

By using different engineered human colorectal cell lines, we first identified RIP140 as a new modulator of RBPJ activity and, consequently, as a novel actor in the regulation of *HES1* gene expression in the intestinal epithelium. RBPJ mediates gene transcription activation or repression depending on the protein complexes that are recruited. In the absence of nuclear NICD, RBPJ represses Notch target genes through the recruitment of corepressor complexes containing proteins such as HDACs, SMRT, NCoR and/or CtBPs (8). The positive regulation by RIP140 which takes place at the transcriptional level and involves the RBPJ binding site, might, at least partly, implicate squelching of HDACs or CtBPs since we previously reported that RIP140 directly interacted with these transcriptional repressors (39). Concerning the induction of *HES1* gene expression by RIP140, we clearly showed that it implicates the RBPJ binding site present in the proximal region of the *HES1* promoter. However, we cannot exclude that RIP140 might activate *HES1* gene transcription through other signaling pathways such as the Wnt, Hedgehog, TGFβ/BMP or hypoxia pathways which can all affect Notch regulation of *HES1* gene expression (49).

Interestingly, **RIP140 inhibits *HES1* gene transcription when the HES1 protein is highly expressed**. In many types of cultured cells including fibroblasts, myoblasts, and neuroblasts, *HES1* exerts a negative feedback loop on its one expression leading to an autonomous oscillatory regulation of its expression with a periodicity of about 2hrs. Hes1 thus acts as a biological clock capable of controlling the activation time of various biological processes such as cell cycle or differentiation (50),(51). Interestingly, in pluripotent stem cells derived from the intern cellular mass of a blastocyst, HES1 oscillatory expression contribute to the multiple differentiation properties of these cells (13).

Our data indicate that **RIP140 mainly acts through the amplification of the repressive activity that HES1 exert on its own promoter** and, as a consequence, participates in the negative feed-back loop that HES1 exerts on its own expression. This effect might be a general one since it was also observed on the *HES4* gene whose expression is also repressed by HES1. Our data obtained using *in vitro* interaction, proximity ligation assays and co-localization identify RIP140 as a new partner of HES1 and further work is needed to fully decipher the underlying mechanisms. It should be noted that little is known about the way HES1 represses gene expression. The C-terminal WRPW domain of the protein interacts with TLE/Grg corepressors (52). On the other hand, the bHLH domain of HES1 recruits Sirt1, a class III histone deacetylase, and thereby represses target gene expression (53).

The molecular machinery involved in the oscillation of *HES1* gene expression is not yet fully understood. It has been reported that the Jak-Stat signaling pathway (54) and the mir-9 (55),(56) could be involved in this regulation. Yet, another mechanism used by HES1 to repress the expression of its target genes is by preventing the activity of transcription activators (50). RIP140 could therefore titrate transcription activators, like ß-catenin, and thus prevent their activity on the *HES1* gene promoter. More importantly, **RIP140 is clearly engaged in a negative regulatory loop due to the induction of its expression by HES1**. Although the HES1 transcription factor has been identified as a repressor of gene expression, it is also able to activate transcription. Several target genes induced by HES1 in CRC cells have been identified including the oncogene *BMI1* (57), which promotes invasion and migration of colon cancer cells.

In link with the numerous cross-talks that occurred between the two transcription factors at different levels, we demonstrated that, in colon cancer cells, **RIP140 is able to reverse the mitogenic effects of HES1** which switched to a repressor of cell proliferation in the presence of ectopic expression of RIP140. Based on these results, it is tempting to speculate that RIP140 might regulate other oncogenic functions of HES1 that have been identified such as its role in metastatic potentiation (58),(59). This hypothesis is sustained by our observation in two cohorts of human colorectal cancer biopsies, in which we observed that **high coexpression of RIP140 and HES1 correlates with a better patient overall survival**.

In conclusion, this study demonstrates for the first time that, in CRC cells, the transcription coregulator RIP140 1) strongly cross-talks and directly interacts with the HES1 transcription factor, 2) is engaged in the autoregulation of its oscillatory expression, 3) reverses its mitogenic activity on cancer cell proliferation and 4) is required to reveal its correlation with prolonged patient survival. This work thus reinforces the role that RIP140 plays in intestinal tumorigenesis by controlling a major molecular pathway involved in colon cancer.

## ACKNOWLEDGEMENTS

We thank the Réseau d’Histologie Expérimentale de Montpellier (RHEM) for histology facilities. We are also grateful to Drs L. Cheng, R Kageyama and M Plateroti for plasmid sharing. We deeply thank A. Boulahtouf for experimental advices and support.

## LIST OF ABBREVIATIONS

APC: Adenomatous polyposis coli
BMI1: 
CRC: colorectal cancer
CtBP: C-terminal binding protein
CtIP: CtBP interacting protein
HES1: hes family bHLH transcription factor 1
HEY: hairy ears, Y-linked
MAML1: Mastermind-like 1
NF-κB: nuclear factor kappa B
NICD: Notch intracellular domain
NRIP1: Nuclear Receptor-Interacting Protein 1
RBPJ: recombination signal binding protein for immunoglobulin kappa J region
RIP140: Receptor Interacting Protein of 140 kDa

