## Supplemental data for "RIP140 regulates HES1 oscillatory expression and mitogenic activity in colon cancer cells"

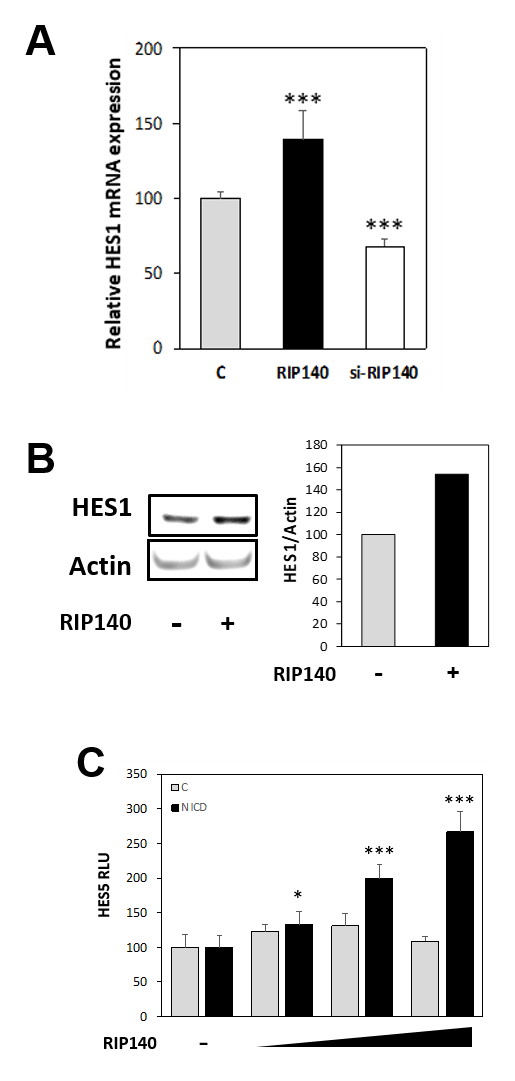
**SUPPLEMENTARY Figures**

**Figure S1: RIP140 increases HES1 expression**

**(A)** *HES1* mRNA level in HT29 CRC cells transiently transfected as in Figure 1A. Results are expressed as fold change ± S.D. relatively to controls; n = 4 to 8 independent experiments for each condition. **(B)** Western blot analysis of HES1 protein level in HT29 cells transiently transfected with the RIP140 expression vector. **(C)** A reporter construct encompassing the *HES5* gene promoter was co-transfected into SW620 cells with increasing doses of RIP140 expression vector in the presence or not of NICD expression vector; n=3 independent experiments.

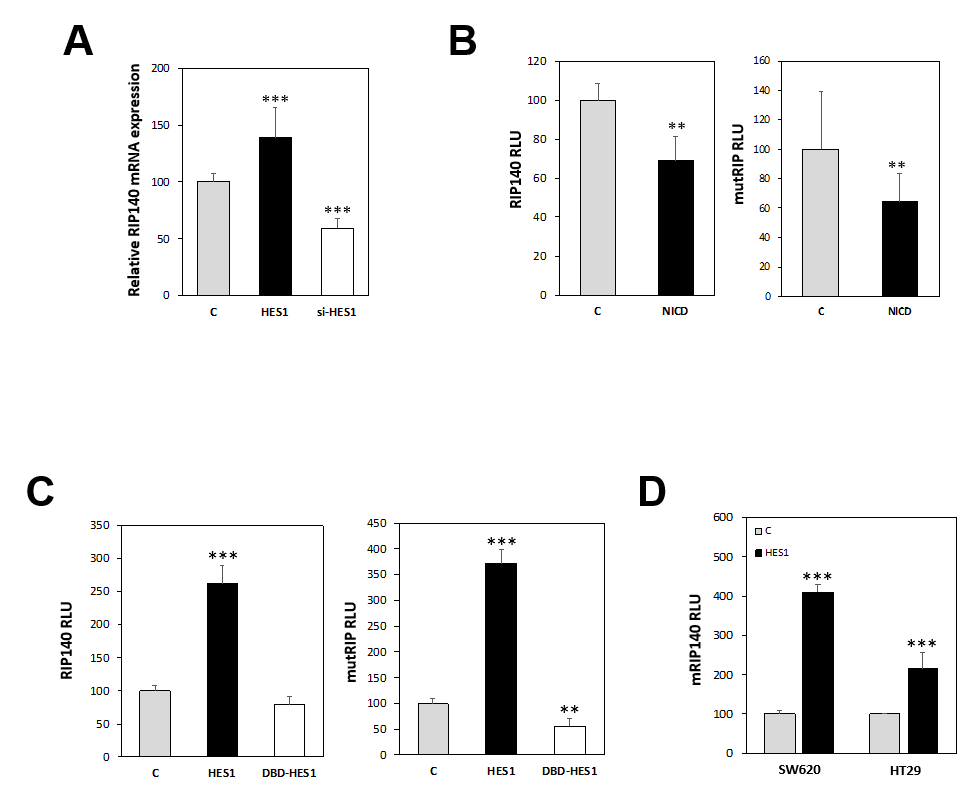
**Figure S2: The *RIP140* gene is a target of the Notch/HES1 pathway**

**(A)** *RIP140* mRNA level in HT29 cells transiently transfected with HES1 expression vector or treated with siRNA targeting the *HES1* mRNA; n = 4 independent experiments. **(B)** Luciferase reporter assay performed on the *RIP140* gene promoter construct (left panel) or on a reporter construct encompassing the very proximal region of *RIP140* gene promoter (right panel) transiently co-transfected into HT29 cells with the NICD expression vector. **(C)** Luciferase reporter assay performed on *RIP140* gene promoter constructs transiently co-transfected into HT29 cells with the HES1 or DBD-HES1 expression vectors; n = 3 independent experiments. **(D)** Luciferase reporter assay performed on a reporter construct encompassing the murine form of RIP140 promoter transiently co-transfected into SW620 or HT29 cells with HES1 expression vector; n = 3 independent experiments.

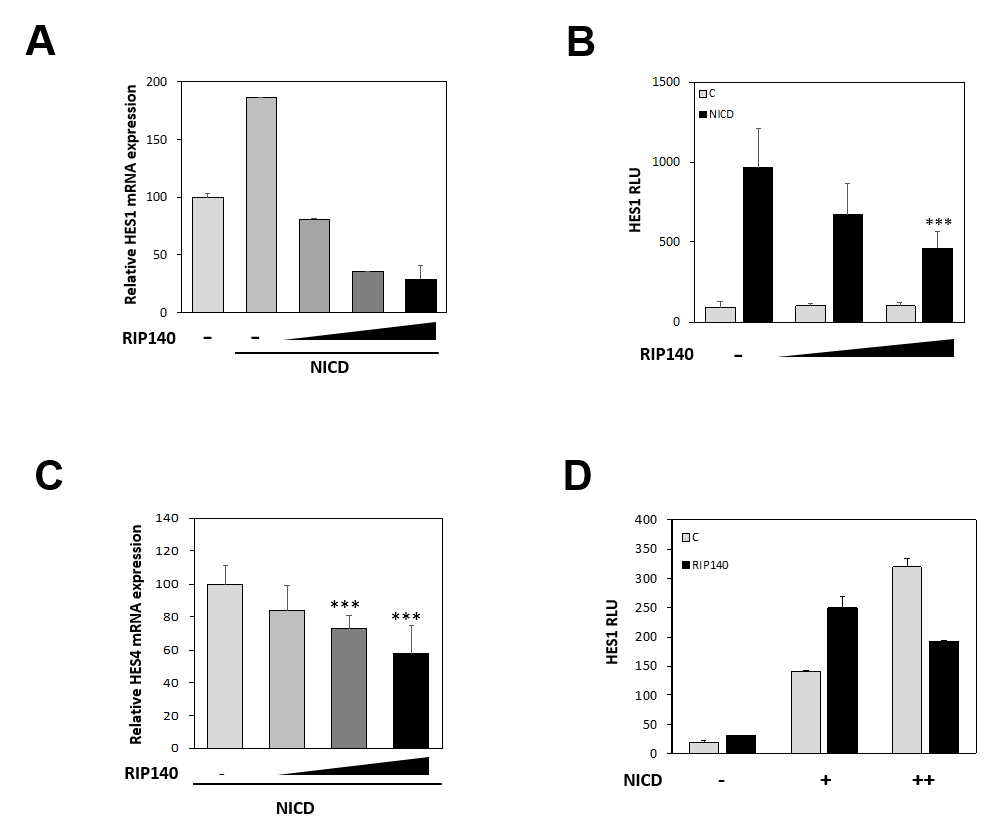
**Figure S3: The *RIP140* gene is a target of the Notch/HES1 pathway**

**(A)** *HES1* mRNA level in HT29 cells transiently transfected with increasing doses of RIP140 expression vector in the presence or not of a high dose of NICD expression vector. **(B)** Luciferase reporter assay performed on *HES1* gene promoter (0.47kb) construct transiently co-transfected into HT29 cells with increasing doses of RIP140 expression vector in the presence or not of the NICD expression vector. **(C)** *HES4* mRNA level in HT29 cells under the same conditions as in panel A. Results are expressed as fold change ± S.D. relatively to control; n = 3 independent experiments. **(D)** Luciferase reporter assay performed on the *HES1* gene promoter (0.47kb) construct transiently co-transfected into HT29 cells with doses of RIP140 and/or NICD expression vectors.

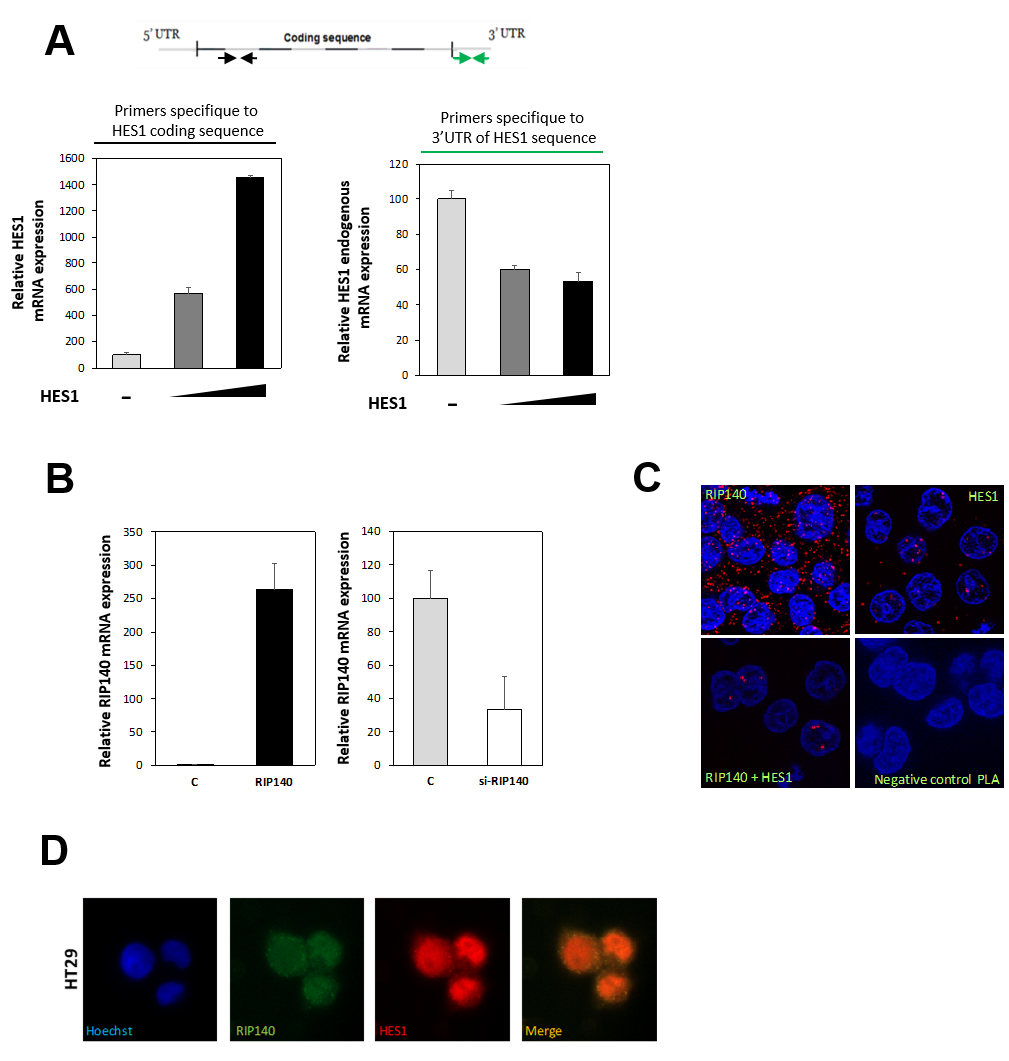
**Figure S4: RIP140 is required for the HES1 feed-back loop**

**(A)** Total (left panel) and endogenous (right panel) *HES1* gene expression distinctly detected using primers specific to the HES1 coding sequence and the 3’UTR of HES1 sequence, respectively in SW620 cells transiently transfected with increasing doses of LV-HES1 expression vector. **(B)** RT-qPCR analysis of *RIP140* mRNA level in SW620 cells transiently transfected with RIP140 expression vector or with the control siRNA (C) or siRNA targeting RIP140 (si-RIP140). **(C)** DuoLink proximity ligation assay performed to visualize endogenous HES1 and RIP140 interaction in HT29 cells. **(D)** Double immunofluorescence analysis (40x) of HES1 and RIP140 protein levels in HT29 cells.

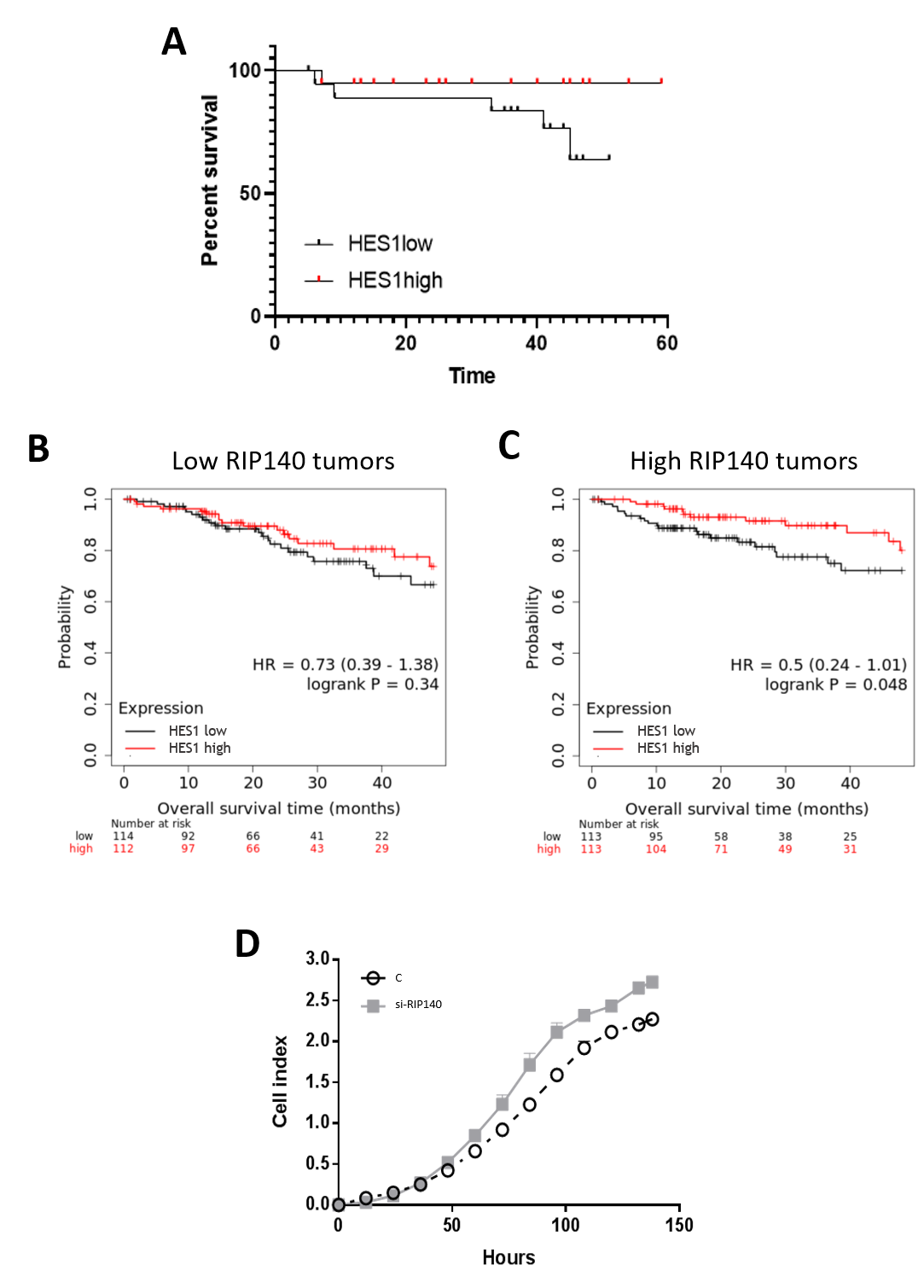
**Figure S5: HES1/RIP140 interplay on intestinal tumorigenesis, CRC cell proliferation and patient survival.**

**(A)** Kaplan-Meier analysis performed on HES1 IHC data. Patients were ranked according to HES1 staining IRS in their tumors and divided into two groups exhibiting low and high expression, respectively (best cut-off threshold). **(B)** and **(C)** Kaplan-Meier analysis of the cumulative OS of patients with low or high HES1 gene expression was performed using the Kaplan–Meier plotter database on the groups exhibiting low (panel B) or high (panel C) RIP140 gene expression. A log-rank test was used for statistical analysis. **(D)** HT29 cell proliferation was measured by xCELLigence assay after transfection or not of a siRNA targeting RIP140.

**Table S1: Primer sequences**

| **Gene** | **Forward Sequence** | **Reverse Sequence** |
| --- | --- | --- |
| **hRIP140** | AATGTGCACTTGAGCCATGATG | TCGGACACTGGTAAGGCAGG |
| **hHES1** | AAGAAAGATAGCTCGCGGCAT | CCAGCACACTTGGGTCTGT |
| **hNotch1** | GAATGGTCAATGCGAGTGGC | GGCCCTGGTAGCTCATCATC |
| **hHES4** | TGGACGCCCTCAGAAAAGAG | TTCACCTCCGCCAGACACT |
| **hHES15’** | GGTGCTGATAACAGCGGAAT | TTGGAGTTCTTCACGAAAAAGA |
| **hp21** | TGAGCGATGGAACTTCGAC | ACAAGACAGTGACAGGTCC |
| **hp27** | AGCCTGGAGCGGATGGACGCC | CTCCCGCTGACATCCTGGCTC |
| **hMuc2** | GACACCATCTACCTCACCCG | TGTAGGCATCGCTCTTCTCA |
| **mRip140** | AGAACGCACATCAGGTGGCA | GATGGCCAGACACCCCTTTG |
| **mHES1** | GCGAAGGGCAAGAATAAATG | TGTCTGCCTTCTCTAGCTTGG |
| **mp21** | GTTCCGCACAGGAGCAAAGT | ACGGCGCAACTGCTCAC |
| **mMuc2** | CGACACCAGGGATTTCGCTTAAT | CACTTCCACCCTCCCGGCAAAC |
| **ComR1** | CCAGTGCTAGCATTCGCTGT | CTAAATCAGAACCCACCCCGGAT |
| **VillinCre** | CAAGCCTGGCTCGACGGCC | CGCGAACATCTTCAGGTTCT |
| **28S** | CGATCCATCATCCGCAATG | AGCCAAGCTCAGCGCAAC |
| **RS9** | CGGCCCGGGAGCTGTTGACG | CTGCTTGCGGACCCTAATGTGACG |
| **ChIP HES1 prom** | GCGTGTCTCCTCCTCCCATT | CCTGGCGGCCTCTATATATA |
